# Nuclear NAD^+^ homeostasis is essential for naive and chemoresistant *BRCA1/2*-deficient tumor survival

**DOI:** 10.1101/2021.12.09.471907

**Authors:** Daniele Musiani, Hatice Yücel, Laura Sourd, Elisabetta Marangoni, Raphael Ceccaldi

## Abstract

Resistance to PARP inhibitors (PARPi) is emerging as the major obstacle to their effectiveness for the treatment of *BRCA1/2*-mutated, also referred as homologous recombination (HR)-deficient, tumors (HRD). Over the years, mechanistic studies gained insights on effectors acting downstream of PARP1, lagging behind the understanding of earlier events upstream - and thus independent - of PARP1. Here, we investigated the role of nuclear NAD^+^, an essential cofactor for the activity of key DNA repair proteins, including PARP1 and sirtuins. We show that NMNAT1-the enzyme synthesizing nuclear NAD^+^ - is synthetically lethal with *BRCA1/2* in a PARP1-independent but SIRT6-dependent manner. Consequently, inhibition of NMNAT1/SIRT6 axis not only kills naive but also PARPi-resistant HRD cancer cells. Our results unravel a unique vulnerability of HRD tumors, therapeutically exploitable even upon PARPi resistance development.

**One-Sentence Summary:** Targeting NMNAT1 kills chemoresistant and naive BRCA1/2-deficient tumors by disrupting SIRT6-dependent base excision repair.

## Results

In the last decade, the synthetic lethality between the poly (ADP-ribose) polymerase-1 (*PARP1)* and *BRCA1/2* (*1*, *2*) has been exploited in the clinic with the approval of PARP inhibitors (PARPi) for the treatment of *BRCA1/2*-mutated tumors (*3*), also referred as homologous recombination (HR)-deficient tumors (HRD) (*4*). More recently, novel vulnerabilities in HRD cells have been identified, thus paving the way for innovative therapies (*5*–*9*). However, resistance to PARPi is emerging as the major obstacle to their clinical effectiveness (*10*–*12*). As a consequence, many patients run out treatment options and succumb to the disease, stressing the need for additional therapeutics (*13*, *14*).

Nicotinamide adenine dinucleotide (NAD+) is extremely compartmentalized in the cell (*15*). While cytosolic and mitochondrial NAD+ pools are essential for global bioenergetic processes such as glycolysis and oxidative phosphorylation, nuclear NAD^+^ acts as an essential cofactor for key DNA repair enzymes, including PARP1 (*16*) and the sirtuin deacetylases (Fig. 1A) (*17*).

**Fig. 1.**
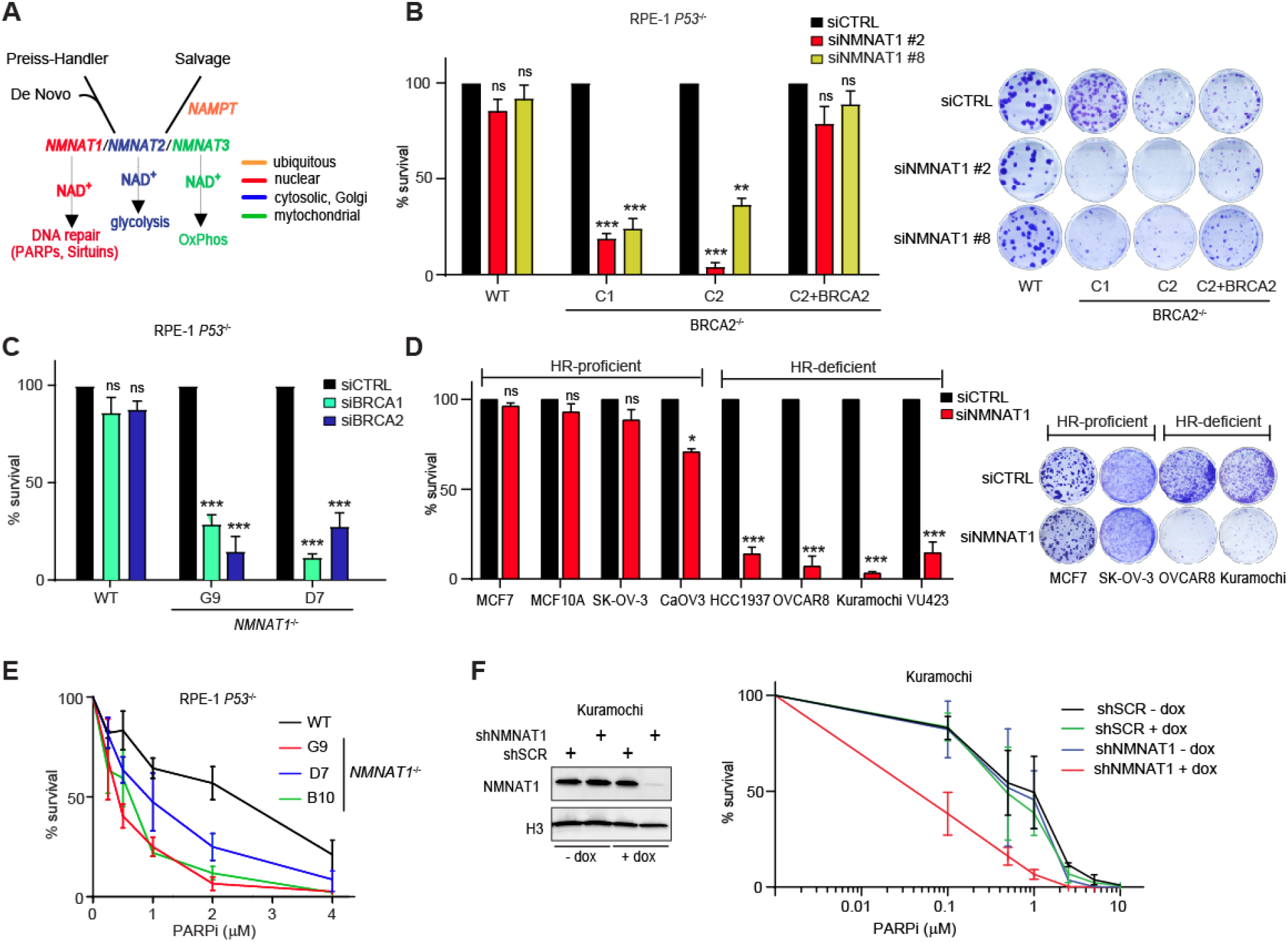
*NMNAT1* is synthetic lethal with *BRCA1/2*. **(A)** Schematic of the compartmentalized NAD^+^ biosynthesis pathways and the different roles of subcellular NAD^+^ pools. (**B)** Quantification (*left*) and representative images (*right*) of clonogenic assay of wild type (WT) and *BRCA2^−/−^* (clones C1 and C2) RPE-1 cells with and without BRCA2 cDNA complementation, after transfection with the indicated NMNAT1 siRNAs. **(C)** Clonogenic formation of WT and *NMNAT1*^−/−^ RPE-1 cells after transfection with the indicated BRCA1/2 siRNAs. **(D)** Quantification (*left*) and representative images (*right*) of clonogenic formation of the indicated HR-proficient and HR-deficient cell lines upon siNMNAT1 #2. **(E and F)** Survival assay of WT-*NMNAT1*^−/−^ RPE-1 **(E)** and Kuramochi cells **(F)** exposed to the indicated doses of the PARP inhibitor (PARPi) rucaparib. Data in **(B-F)** represent mean ± s.e.m. *n* ≥ 3 (independent experiments); ns, non significant; *, *p* < 0.05; **, *p* < 0.01; ***, *p* < 0.001 (two-tailed *t* test).

Given the reliance of HRD cells on PARP1 activity (*1*, *2*), we reasoned that nuclear NAD^+^ homeostasis is a critical determinant of HRD cell survival and thus nuclear NAD^+^ levels must be finely tuned in those cells. For this reason, we explored the role of nicotinamide nucleotide adenylyl transferase 1 (NMNAT1), the sole enzyme catalyzing the final step of nuclear NAD^+^ synthesis (*18*, *19*), in the maintenance of genome stability and survival of HRD tumors.

### NMNAT1 is synthetically lethal with BRCA1/2

We generated various HRD isogenic cell systems using CRISPR-Cas9 mediated genome editing. We knocked-out *BRCA2* gene in *TP53*^−/−^ epithelial RPE-1 cells and knocked-in a mini auxin-inducible degron (mAID) at *BRCA2* locus in HeLa cells. In those isogenic models, HR was compromised by genetic deletion in *BRCA2*^−/−^ RPE-1 clones (Fig. S1, A and B), or by auxin (IAA) treatment in mAID *BRCA2* HeLa cells (Fig. S1C), as shown by impairment of ionizing radiation-induced RAD51 foci formation (Fig. S1, D-F) and PARPi hypersensitivity (Fig. S1, G and H).

In RPE-1 cells, *NMNAT1* knockdown by two different short interphering RNA sequences (siRNA) (Fig. S1I) impaired the clonogenic cell survival of the two *BRCA2*^−/−^ clones (C1 and C2), while having no effect on the survival of parental, HR-proficient (HRP) cells (Fig. 1B). Complementation of *BRCA2*^−/−^ cells with a full length BRCA2 cDNA (C2+BRCA2) (Fig. S1B) rescued not only PARPi hypersensitivity (Fig. S1G), but also cell survival upon *NMNAT1* depletion (Fig. 1B). Likewise, in the mAID *BRCA2* HeLa isogenic model, *NMNAT1* knockdown by short hairpin RNA (shRNA) reduced clonogenic survival only in the presence of auxin, *i.e.* upon BRCA2 protein degradation (Fig. S1J). Furthermore, NMNAT1 silencing also impaired the clonogenic ability of the previously generated *BRCA1* knock-out (*BRCA1*^−/−^) *TP53*^−/−^ RPE-1 cells (Fig. S1, B-E), while sparing parental HRP cells (Fig. S1K). Altogether, these data indicate that *NMNAT1* is synthetically lethal with *BRCA1/2*.

To corroborate these findings, we used a complementary approach. We generated several *NMNAT1* knockout (*NMNAT1*^−/−^) clones in *TP53*^−/−^ RPE-1 cells (Fig. S2A) and assessed genome integrity and survival upon *BRCA1*/*2* depletion. When compared to parental cells, *NMNAT1*^−/−^ clones showed a mild defect in proliferation (Fig. S2B) and a slight increase in DNA breaks formation, as measured by both comet assay (Fig. S2C) and γH2AX immunofluorescence (Fig. S2D). However, siRNA-mediated *BRCA1/2* depletion (Fig. S2E) severely impaired the clonogenic survival of the *NMNAT1*^−/−^ clones (Fig. 1C), while having no significant effect on parental cell survival further confirming a synthetic lethal interaction between *NMNAT1* and *BRCA1/2*.

Next, we assessed the survival ability of a panel of HRP and HRD cancer cells upon siRNA-mediated *NMNAT1* depletion. Knockdown of *NMNAT1* abolished the clonogenic ability of HRD cells and induced apoptotic cell death, whereas it had little to no effects in HRP cells, including the immortalized normal breast epithelial MCF-10A cells (Fig. 1D; Fig. S2, F and G).

Altogether, we collected data showing that *NMNAT1* is required for the survival of HRD tumors.

To translate synthetic lethal interactions into cancer treatments, one important consideration is the toxic impact of the inhibition of the gene under investigation in normal tissue (13). To rule out any possible detrimental effect of *NMNAT1* loss in normal cells, we generated several *NMNAT1*^−/−^ clones in *TP53*^+/+^ RPE-1 cells (Fig. S2H) and assessed proliferation and genome integrity, as quantified by DNA breaks measurement by alkaline comet assay. We found that *NMNAT1* loss only slightly affected genome integrity without any impact on cell proliferation (Fig. S2, I and J), suggesting that targeting NMNAT1 could have no toxic impact in normal tissue.

### NMNAT1 promotes cell survival independently of PARP1

NMNAT1 fuels PARP1 activity through nuclear NAD^+^ synthesis (*20*). To understand whether *NMNAT1* is epistatic with *PARP1* in cell survival, we exposed *NMNAT1*^−/−^ and parental RPE-1 cells to the PARP inhibitor rucaparib (PARPi). We found that all the *NMNAT1*^−/−^ clones were sensitized to rucaparib when compared to parental cells (Fig. 1E). To test whether this was also the case in HRD cells, we used the *BRCA2*-mutated ovarian cancer cells Kuramochi expressing a doxycycline-inducible *NMNAT1* shRNA. Similarly, we found that depletion of *NMNAT1* by addition of doxycycline further sensitizes the cells to PARPi (Fig. 1F). Altogether, our data suggest that *NMNAT1* is not fully epistatic with *PARP1* in cell survival, but rather acts through other downstream effectors.

Besides providing fuel for NAD^+^-dependent nuclear enzymes through its catalytic activity, NMNAT1 can also act as a chaperone (*21*). To understand whether the catalytic activity of NMNAT1 is crucial in HRD cells, we complemented *NMNAT1^−/−^* RPE-1 cells with either wild type (WT) or a catalytically-dead version of the enzyme (W169A) (*20*) and tested cell survival upon exposure to PARPi or following *BRCA2* knockdown. We first confirmed the ability of WT NMNAT1, but not W169A mutant, to sustain PARP1 hyper-activation upon methyl methanesulfonate (MMS) exposure (Fig. 2A). Next, we found that the ectopic expression of WT NMNAT1, but not the W169A mutant, reduced the amount of DNA breaks in *NMNAT1*^−/−^ cells back to parental cells levels (Fig. 2B), indicating that the catalytic activity of NMNAT1 is important for genome integrity. Finally, both PARPi hyper-sensitivity and synthetic lethality with *BRCA2* were overcome in *NMNAT1*^−/−^ cells by expressing WT NMNAT1, but not the W169A mutant (Fig. 2, C and D; Fig. S3A). Altogether, these data indicate that the catalytic activity of NMNAT1, i.e. nuclear NAD^+^ homeostasis, is essential to maintain genome integrity and HRD cell survival. Since the catalytic activity of NMNAT1 is not epistatic with PARP1 activity in cell survival, these results also suggest that nuclear NAD^+^-dependent enzymes downstream of NMNAT1, other than PARP1, are crucial for these functions.

**Fig. 2.**
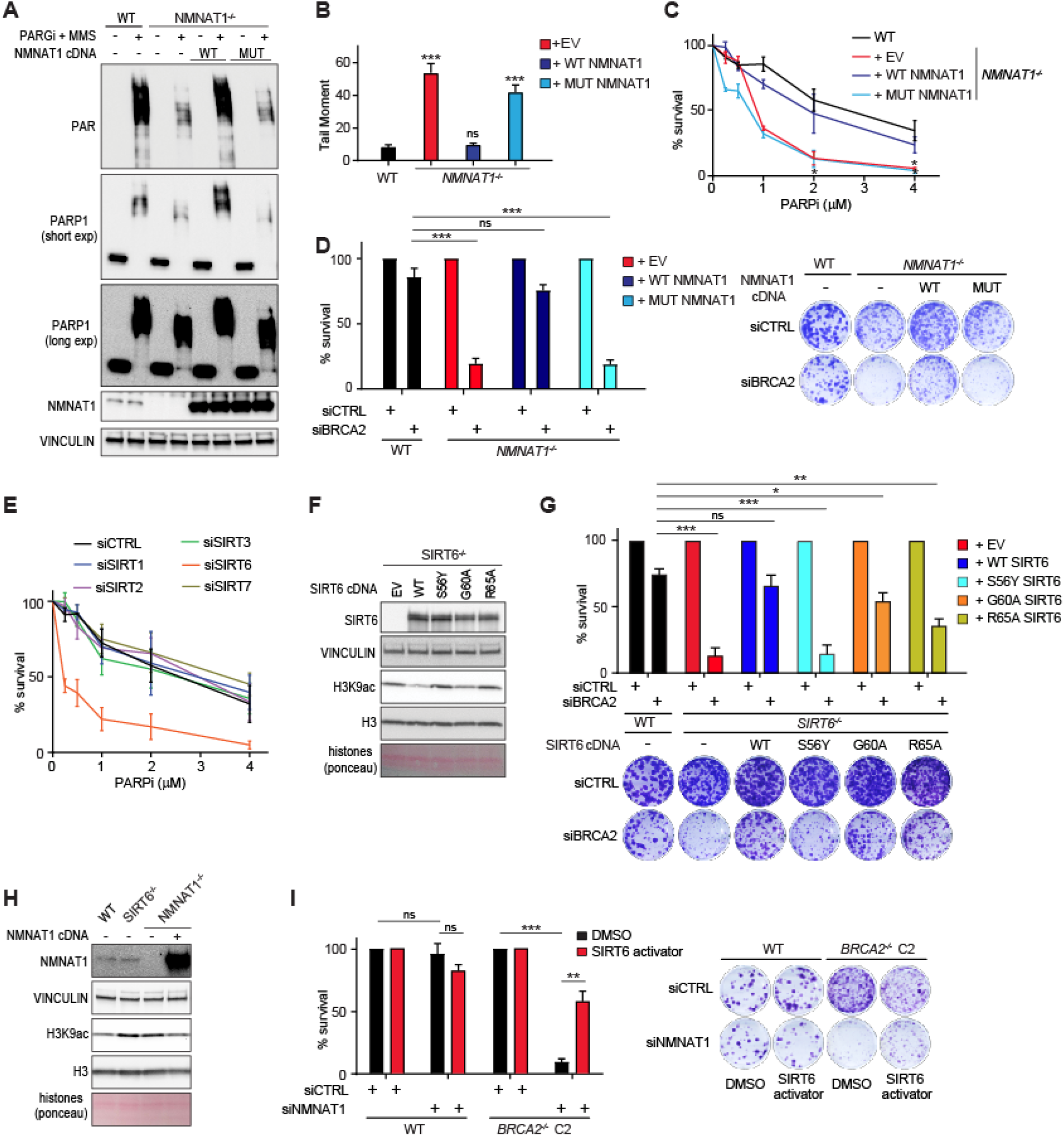
SIRT6 functions downstream of NMNAT1 in the survival of HRD cells. **(A)** Immunoblot analysis showing PARP1 activation in WT and *NMNAT1*^−/−^ RPE-1 cells (G9) complemented with wild type (WT), catalytically dead (MUT, W169A) NMNAT1 cDNA or empty vector (EV) following exposure to 10 mM of methyl methanesulfonate (MMS) for 10 minutes (min) after 2 hours of pre-treatment with 10 μM of PARGi. (**B)** DNA breaks quantification by alkaline COMET assay of WT and *NMNAT1*^−/−^ (clone G9) RPE-1 cells complemented or not with wild type (WT) or catalytically dead (MUT, W169A) NMNAT1 cDNA. **(C and D)** Clonogenic formation of cells as in (**B)** in response to rucaparib (PARPi) **(C)** or following transfection with either *BRCA2* or control (siCTRL) siRNAs **(D***left panel*). Representative images are shown in **(D** *right panel*). **(E)** Survival assay of RPE-1 cells in response to rucaparib (PARPi) after transfection with siRNAs targeting the indicated sirtuins. **(F)** Immunoblot analysis of H3K9ac in *SIRT6*−/− RPE-1 cells complemented or not (EV) with the indicated SIRT6 cDNAs. **(G)** Quantification of clonogenic assay (*upper panel*) and representative images (*lower panel*) of RPE-1 cells as in **(F)** after transfection with si*BRCA2* or control (siCTRL). **(H)** Immunoblot analysis of H3K9ac in *SIRT6*−/− and *NMNAT1−/−* RPE-1 cells, complemented or not with WT NMNAT1 cDNA. **(I)** Clonogenic formation of wild type (WT) and *BRCA2*^−/−^ (C2, clone 2) RPE-1 cells following transfection with si*NMNAT1* or siCTRL after 4 days of pre-treatment with 10 μM of the SIRT6 activator MDL-800. MDL800 was refreshed every 3 days for the duration of the assay. All data represent mean ± s.e.m. *n* ≥ 3 (independent experiments); ns, non significant; *, *p* < 0.05; **, *p* < 0.01; ***, *p* < 0.001 (two-tailed *t* test).

### SIRT6 promotes cell survival independently of PARP1

Besides PARP1, nuclear NAD^+^ is also used as a cofactor by sirtuins, a class of deacetylases involved in many processes, including DNA repair (*17*). Thus, we investigated whether the depletion of any nuclear sirtuin would result in PARPi hypersensitivity, as we observed upon *NMNAT1* loss (Fig. S3B). siRNA-mediated knockdown of *SIRT6*, but not *SIRT1*, *SIRT2*, *SIRT3* or *SIRT7*, sensitized cells to PARPi (Fig. 2E), suggesting that among all the nuclear sirtuins SIRT6 is the key player downstream of NMNAT1 in the response to PARPi. These data also suggest that *SIRT6*, similarly to *NMNAT1*, is not epistatic with *PARP1* in cell survival.

Nonetheless, SIRT6 has been shown to promote DNA repair by activating PARP1 (*22*). Hence, to further investigate the interplay between SIRT6 and PARP1, we generated a *SIRT6* knockout (*SIRT6*^−/−^) clone in *TP53*^−/−^ RPE-1 cells and exposed it to rucaparib. Likewise, *SIRT6*^−/−^ cells were more sensitive than parental cells to PARPi, in a manner that was reminiscent of *NMNAT1*^−/−^ RPE-1 cells (Fig. S3C), confirming that *SIRT6* is not epistatic with *PARP1* in cell survival.

Similar to PARP1, SIRT6 has been shown to function as a DNA double strand breaks (DSBs) sensor (*23*, *24*). In addition, SIRT6 maintains genome stability by promoting various DNA repair pathways, including HR, non-homologous end-joining (NHEJ) and base-excision repair (*25*–*28*). In accordance with these findings, we found that *SIRT6*^−/−^ cells had higher level of DNA breaks than parental cells and that complementation of *SIRT6*^−/−^ cells with WT SIRT6 cDNA rescued DNA breaks accumulation (Fig. S3D).

We next evaluated a potential synthetic lethality between *SIRT6* and *BRCA2,* also in light of a recent report showing an essential role of sirtuins in *BRCA*-deficient cell survival (*29*). Knockdown of *BRCA2* by siRNA severely impaired the survival of *SIRT6*^−/−^ cells, while having no significant effect on parental cell survival (Fig. S3E). Furthermore, SIRT6 inhibition by the selective small molecule compound OSS128167 (SIRT6i) (*30*) killed the two *BRCA2^−/−^* RPE-1 clones (C1 and C2) while sparing parental HRP cells (Fig. S3F). These data show that *SIRT6*, like *NMNAT1*, is synthetically lethal with *BRCA2*.

SIRT6 is a NAD^+^-dependent nuclear enzyme that can deacetylate and mono-ADP ribosylate (MARylate) a variety of target proteins, including histone H3K9 (*31*) and PARP1 (*22*) respectively. To determine which catalytic activity of SIRT6 is important for HRD cell survival, we complemented *SIRT6*^−/−^ cells with either WT or dissociation-of-function SIRT6 mutants, which were previously described (Fig. 2F) (*32*). While the S56Y mutant, which lacks both deacetylation and MARylation activities, did not rescue either PARPi sensitivity or *BRCA2* synthetic lethality, the MAR-dead mutant G60A and the deacetylase-dead mutant R65A did partially rescued both phenotypes, although to a lesser extent than the WT SIRT6 (Fig. S3G; Fig. 2G). Together, these data indicate that both of the NAD^+^-dependent catalytic activities of SIRT6 play a role in the cellular response to PARPi and in the survival of HRD cells.

Next, we hypothesized that SIRT6 could function downstream of NMNAT1 in the maintenance of genome integrity and HRD cell survival. As a matter of fact, SIRT6 activity was decreased in *NMNAT1*^−/−^ cells, as shown by increased acetylation of H3K9 (H3K9ac) as compared to parental cells (Fig. 2H). We found that the SIRT6 activator MDL-800 (*33*), which was shown to boost SIRT6 deacetylase function reduced the DNA breaks observed in *NMNAT1*^−/−^ cells to a level similar than in WT parental cells (Fig. S3H). Furthermore, MDL800 pre-treatment also rescued the survival of *BRCA2*^−/−^ cells following *NMNAT1* knockdown (Fig. 2J). These results identify SIRT6 as the key player downstream of NMNAT1 in maintaining genome stability and HRD cell survival. To corroborate this concept, we generated *NMNAT1*^−/−^/SIRT6^−/−^ cells (Fig. S3I) and tested their response to *BRCA2* knockdown. The double-knockout cells were sensitive to *BRCA2* depletion to the same extent as the single knockout cells (Fig. S3J), confirming that NMNAT1 and SIRT6 act epistatically in maintaining the viability of *BRCA*-deficient cells.

### The NMNAT1/SIRT6 axis is required for PARPi-resistant HRD cell survival

Despite the striking cytotoxicity of PARPi in *BRCA*-mutated cells, insurgence of resistance is ubiquitous in the clinic and calls for the design of alternative therapies for the treatment of advanced diseases (*34*). Our findings that inhibition of NMNAT1/SIRT6 kills HRD cells in a PARP-independent manner suggest that targeting this signaling axis might also kill PARPi-resistant HRD cells. To test this hypothesis, we took advantage of preexisting models, such as cell lines derived from patients, as well as generated several cellular models with various mechanisms of PARPi resistance, including fork stabilization and HR restoration (*35*) (Fig. S4A).

First, we generated resistant cells by continuous exposure of *BRCA2*^−/−^ RPE-1 to rucaparib (Fig. S4B). We did not observe HR restoration in any of the derived clones, but resistance likely arose through replication fork stabilization, as shown by the absence of IR-induced RAD51 foci formation and the results of the combing assay in C2-1 and C2-3 clones (Fig. S4, C and D). Interestingly, *NMNAT1* knockdown by siRNA or the use of SIRT6i impaired the clonogenic ability of these PARPi-resistant clones to a similar extent to that of the drug-naive *BRCA2*^−/−^ cells, while sparing the HRP RPE-1 (Fig. 3A). In addition, several clones were derived from the *BRCA2*-mutated pancreatic cancer cell line CAPAN-1 as well, in a similar manner to RPE-1 cells (Fig. S4E). These clones did not restore HR either (Fig. S4F) but rather developed PARPi resistance through other mechanisms. Likewise, shRNA-mediated knockdown of *NMNAT1* or *SIRT6* (Fig. S4G) impaired the clonogenic survival of all the CAPAN-1-derived resistant clones, while having no effect on the HRP normal breast epithelial cell line MCF10A (Fig. 3B). Together, these data show that the synthetic lethality between *NMNAT1* or *SIRT6* and *BRCA1/2* is maintained in PARPi-resistant HRD cells.

**Fig. 3.**
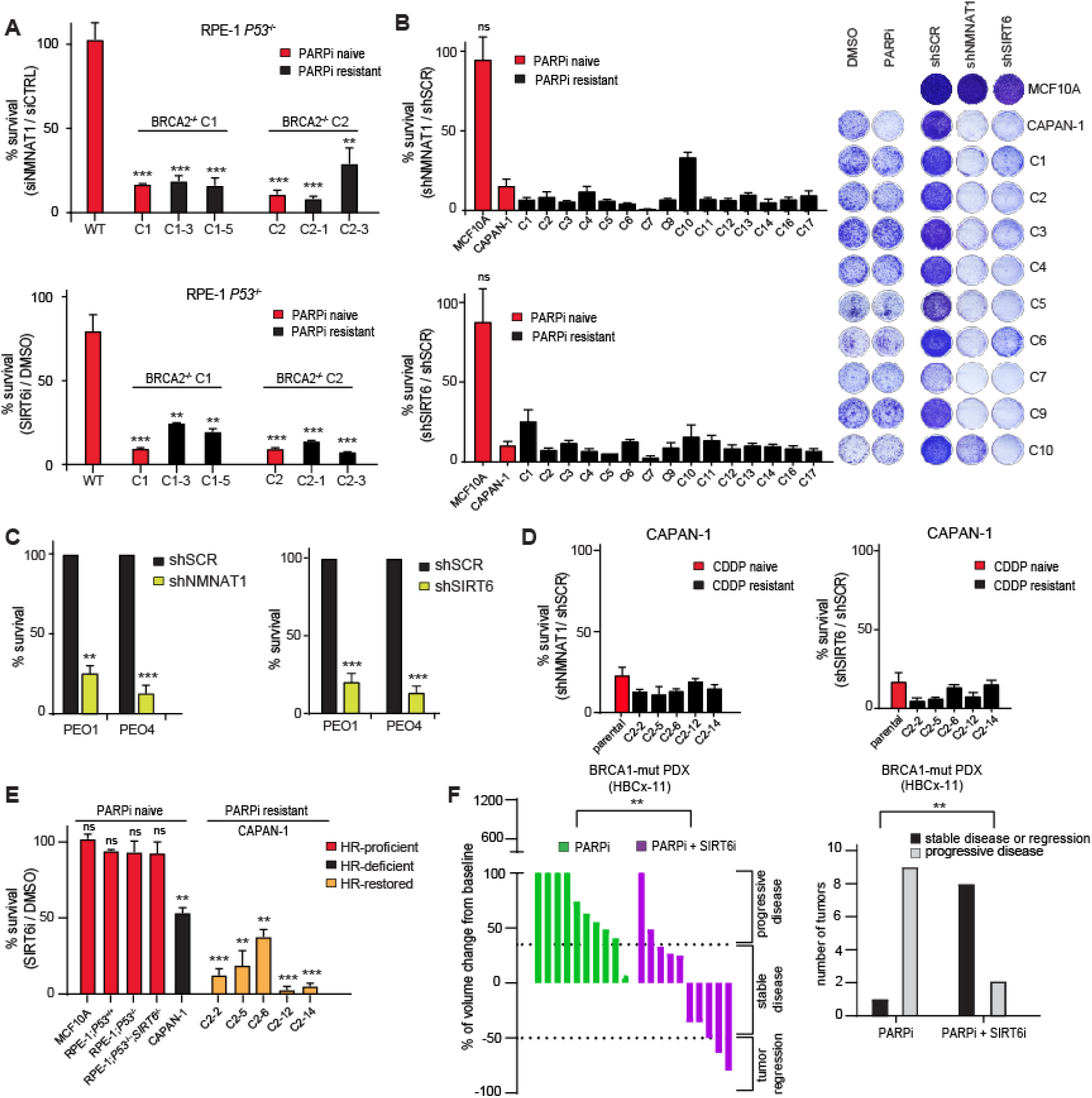
Inhibition of NMNAT1 or SIRT6 kills PARPi-resistant HRD cells, regardless the mechanism of resistance. **(A)** Clonogenic formation of wild type (WT), *BRCA2^−/−^* (clones C1 and C2) and *BRCA2^−/−^*-derived PARPi-resistant RPE-1 cells after control or NMNAT1 (siNMNAT1 #2) siRNAs transfection (*upper panel*) or following exposure to 0.5 mM of the SIRT6 inhibitor OSS_128167 (SIRT6i) (*lower panel*). (**B)** Clonogenic formation of parental and derived PARPi-resistant CAPAN-1 cells after transduction with shNMNAT1 (*upper left panel*) or shSIRT6 (*lower left panel*) lentiviral particles. The HR-proficient normal cell line MCF10A was used as a control. Reprentative images of a selection of cell lines are shown in the *right panel*. **(C and D)** Clonogenic formation of (**C)** PEO-1 and PEO-4 and (**D)** parental and derived cisplatin-resistant CAPAN-1 cells after transduction with shNMNAT1 (*left panels*) or shSIRT6 (*right panels*) lentiviral particles. Lentiviral particles carrying control shRNA (shSCR) were used as control for the experiments shown in **(B-D)**. **(E)** Clonogenic formation of the indicated cell lines following continuous exposure to 0.5 mM SIRT6 inhibitor OSS_128167 (SIRT6i). **(F)** Waterfall plot (*left panel*) of the PARPi-resistant BRCA1-mutated PDX HBCx-11 at day 32 upon treatment with either the PARP inhibitor olaparib (PARPi) or in combination with the SIRT6 inhibitor OSS_128167 (SIRT6i). The number of tumors classified either as stable or progressive disease are plotted on the *right panel*. Data in **(A-E)** represent mean ± s.e.m. *n* ≥ 3 (independent experiments); ns, non significant; *, *p* < 0.05; **, *p* < 0.01; ***, *p* < 0.001 (two-tailed *t* test). In **(B and D)**, all the samples have ***, *p* < 0.001 with the exception of C10 in (**B** *upper panel*)(**, *p* < 0.01), unless otherwise stated.

Nonetheless, HR restoration by secondary mutations in the *BRCA2* gene is the only mechanism of resistance to PARPi which has been observed in patients so far (*10*, *11*). Therefore, we evaluated the effect of NMNAT1/SIRT6 inhibition in HRD cells that developed chemo-resistance through *BRCA2* secondary mutations restoring the open reading frame of the gene and thus HR. In particular, we tested the chemo-resistant HR-restored ovarian cancer cell line PEO4 together with the paired parental *BRCA2*-mutated PEO1 and five clones derived from prolonged *in vitro* cisplatin treatment of CAPAN-1 cells, each bearing different secondary mutation in BRCA2 (*11*, *35*). Surprisingly, shRNA-mediated depletion of *NMNAT1* or *SIRT6* impaired the survival ability of PEO4 and all the resistant CAPAN-1 clones to an extent similar to what we observed in naive cells (Fig. 3, C and D). In addition, SIRT6i killed all the resistant CAPAN-1 clones, while sparing HRP cells including MCF10A (Fig. 3E). Finally, to translate our findings into a translational setting, we used the patient-derived xenograft (PDX) HBCx-11 model, established from a BRCA1-mutated triple negative breast cancer (TNBC) resistant to PARPi. The HBCx-11 tumor was transplanted in nude mice, which were further separated in groups receiving either vehicle, PARPi or the combination of PARPi + SIRT6i. While PARPi alone had almost no effect on tumor regression (1 tumor out of 10 showed a response), we found that the addition of the SIRT6i to the chemotherapy dramatically enhanced the tumor response (8 tumors out of 10) (Fig. 3F).

Altogether our data show that targeting NMNAT1/SIRT6 kills chemo-resistant HRD cells regardless the mechanism of acquired resistance.

### NMNAT1/SIRT6 promotes survival of BRCA1/2-deficient cells by sustaining base excision repair

Genomic instability is often due to defects in the DNA repair machinery and accompanied by increased sensitivity to genotoxic drugs. To understand in which DNA repair pathway NMNAT1/SIRT6 functions, *NMNAT1^−/−^*, SIRT6^−/−^ and parental RPE-1 cells were exposed to various DNA damaging agents and cell survival was evaluated. We found that both *NMNAT1* and *SIRT6* knockout cells were not only more sensitive to MMS than parental cells but also slightly sensitized to IR (Fig. 4, A and B). This is reminiscent of a defect in base excision DNA repair (BER). To evaluate whether the role of NMNAT1/SIRT6 in BER is essential for *BRCA1/2*-deficient cell survival, we over-expressed in our cellular models the lyase domain (8 kDa) of Polβ (*POLB*) (Fig. S5 A-E), which is essential to catalyze the rate-limiting step in BER (*36*, *37*). Then, we evaluated its ability to rescue the synthetic lethality between *NMNAT1/SIRT6* and *BRCA1/2*. We observed that the cytotoxicity of *BRCA2* knockdown in *NMNAT1* or *SIRT6* RPE-1 knockout cells was overcome by over-expression of the 8kDa Polβ (Fig. 4, C and D), suggesting that the BER defects imposed by the lack of NMNAT1/SIRT6 are deleterious for *BRCA2*-deficient cells. Likewise, the sensitivity of the HR-deficient cancer cell line OVCAR8 to *NMNAT1* knockdown was partially rescued by the 8kDa Polβ over-expression (Fig. 4E). In addition, the BRCA2-mutated CAPAN-1 cells and the *BRCA2^−/−^* RPE-1 clone C1 were both partially protected from the SIRT6i cytotoxicity upon 8kDa Polβ over-expression (Fig. 4F). These data indicate that NMNAT1/SIRT6 axis promotes the survival of BRCA1/2-deficient cells by sustaining BER.

**Fig. 4.**
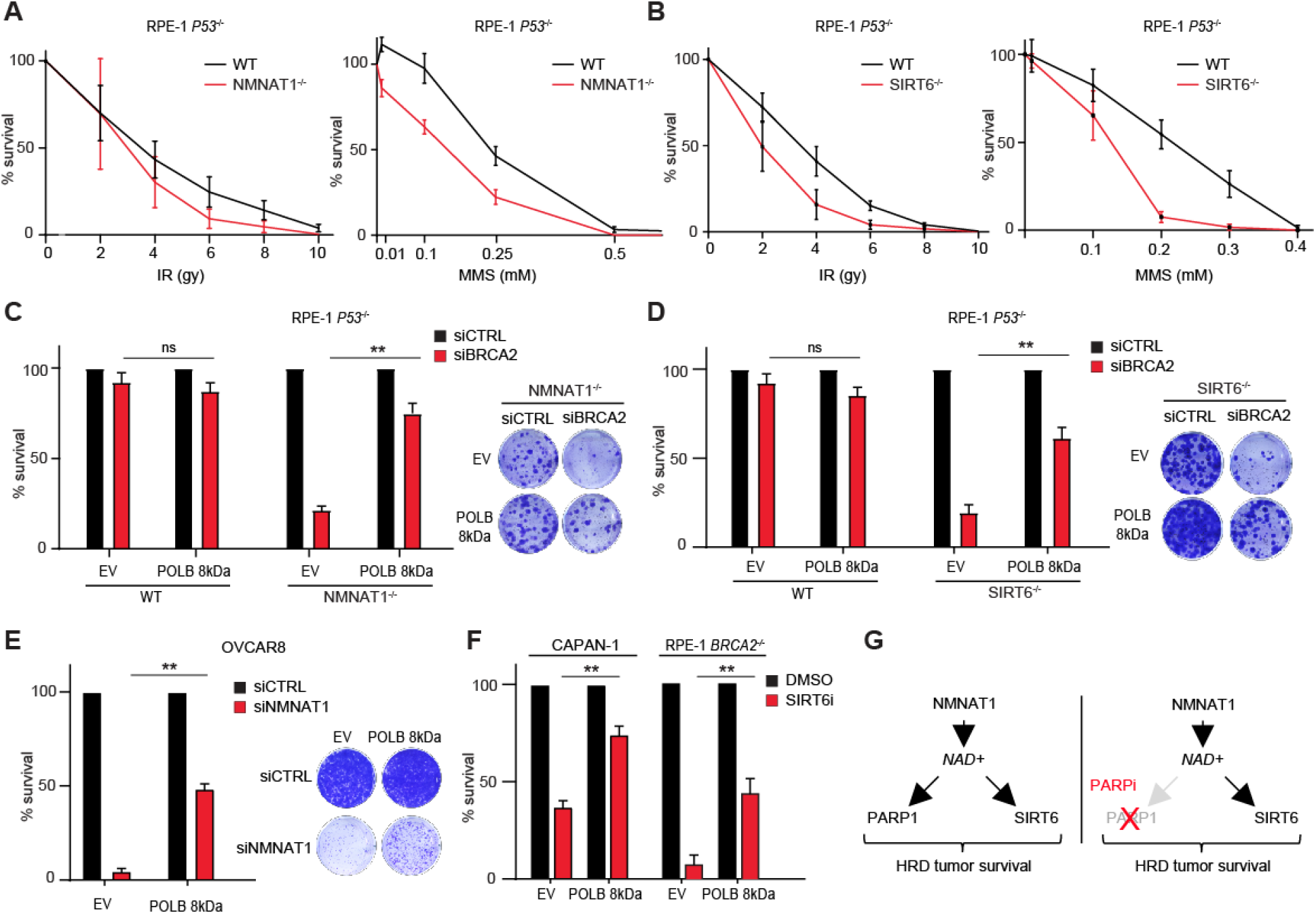
Over-expression of POLB lyase domain overcomes cytotoxicity of NMNAT1 or SIRT6 inhibition in HRD cells. **(A and B)** Clonogenic formation of wild type (WT) **(A and B)**, *NMNAT1^−/−^* (G9) **(A)** and SIRT6^−/−^ (**B)** RPE-1 cells following exposure to the indicated doses of ionizing radiation (IR) or methyl methanesulfonate (MMS). **(C and D)** Clonogenic formation (*left panels*) and representative images (*right panels*) of empty vector (EV) **(C and D)**, *NMNAT1^−/−^* (G9) (**C)** and SIRT6^−/−^ (**D)** RPE-1 cells with and without over-expression of POLB lyase domain (POLB 8kDa) after control (siCTRL) or BRCA2 (siBRCA2) siRNAs transfection. **(E)** Clonogenic formation (*left panel*) and representative images (*right panel*) of empty vector (EV) and POLB-overexpressing (POLB 8kDa) OVCAR8 cells upon control (siCTRL) or NMNAT1 (siNMNAT1 #2) siRNAs transfection. **(F)** Clonogenic formation (*left panel*) and representative images (*right panel*) of empty vector (EV) and POLB-overexpressing (POLB 8kDa) CAPAN-1 cells following exposure to 0.5 mM SIRT6 inhibitor OSS_128167 (SIRT6i). **(G)** model. All the data represent mean ± s.e.m. *n* ≥ 3 (independent experiments); ns, non significant; *, *p* < 0.05; **, *p* < 0.01 (two-tailed *t* test).

Altogether, our results uncover nuclear NAD^+^ homeostasis as an essential determinant of HRD tumor survival. Interestingly, targeting intracellular NAD^+^ has been thoroughly investigated as an anticancer strategy in previous works, notably through the development of inhibitors of NAMPT (nicotinamide phosphoribosyl transferase), the rate-limiting enzyme in NAD^+^ biosynthesis in mammals (*38*). However, NMNAT1 and NAMPT have a different impact on NAD^+^ biogenesis. While NMNAT1 inhibition only affects nuclear NAD^+^ levels and consequently nuclear NAD^+^-dependent enzymes, NAMPT inhibition has been shown to induce a massive depletion of all intracellular NAD^+^ pools, i.e. nuclear, cytosolic and mitochondrial, thus affecting key cellular pathways of energy metabolism, including glycolysis, tricarboxylic acid (TCA) cycle and oxidative phosphorylation. As a consequence, these effects are likely responsible for the NAMPT inhibitors toxicity observed in clinical trials for cancer, with no compounds reported to have progressed through later stages (*39*). Conversely, our data suggest that finely tuning nuclear NAD^+^ homeostasis can be detrimental to HRD tumor survival without causing toxicity in the normal, non-tumoral cellular compartment.

To conclude, we uncovered the first parallel-to-PARP1 pathway mediating HRD tumor survival. Targeting NMNAT1 kills naive and chemoresistant BRCA1/2-deficient tumors by disrupting SIRT6-dependent base excision repair. Importantly, it also kills HRD tumors with somatic reversion of the BRCA1/2 mutations and HR restoration, the only PARPi- and platinum-resistance mechanism described in patients so far, and for which no therapy is currently available. Future NMNAT1 or SIRT6 inhibitors potentially represent a new class of innovative treatments for chemoresistant HRD tumors (Fig. 4G).

## Acknowledgments

We are grateful to Dr. Toshi Taniguchi (Tokai University School of Medicine, Isehara, Japan) for providing the cisplatin-resistant CAPAN-1 clones and to Alan D. D’Andrea (Dana-Farber Cancer Institute, Boston, USA) for the BRCA1−/− RPE-1 cells. We would like to thank Fariba Nemati, Rania El Botty, Léa Huguet, Ahmed Dahmani for technical help in conducting mice experiments. Finally, we thank Léonie Havé and Marie Regairaz for running DNA comet assays, Damiano Borrello for generating NMNAT1/SIRT6 double-knockout cells, and all members of the Ceccaldi lab for helpful discussions.

## Funding

ERC St Grant (N°714162) (RC)

Emergence Program from the city of Paris (DAE137) (RC)

Site de Recherche Intégrée sur le Cancer (SIRIC) grant (RC).

## Author contributions

HY, DM performed and analyzed all experiments but the Fig. 3F.

LS, EM performed and analyzed the experiment in Fig. 3F.

DM and RC conceived, supervised the study and wrote the manuscript.

## Competing interests

Authors declare that they have no competing interests.

## Data and materials availability

All data are available in the main text or the supplementary materials.

## Supplementary Materials

Figs. S1 to S5

## Notes

### Competing Interest Statement

The authors have declared no competing interest.

