## Supplemental Material for "Nuclear NAD^+^ homeostasis is essential for naive and chemoresistant *BRCA1/2*-deficient tumor survival"

**Fig. S1.**

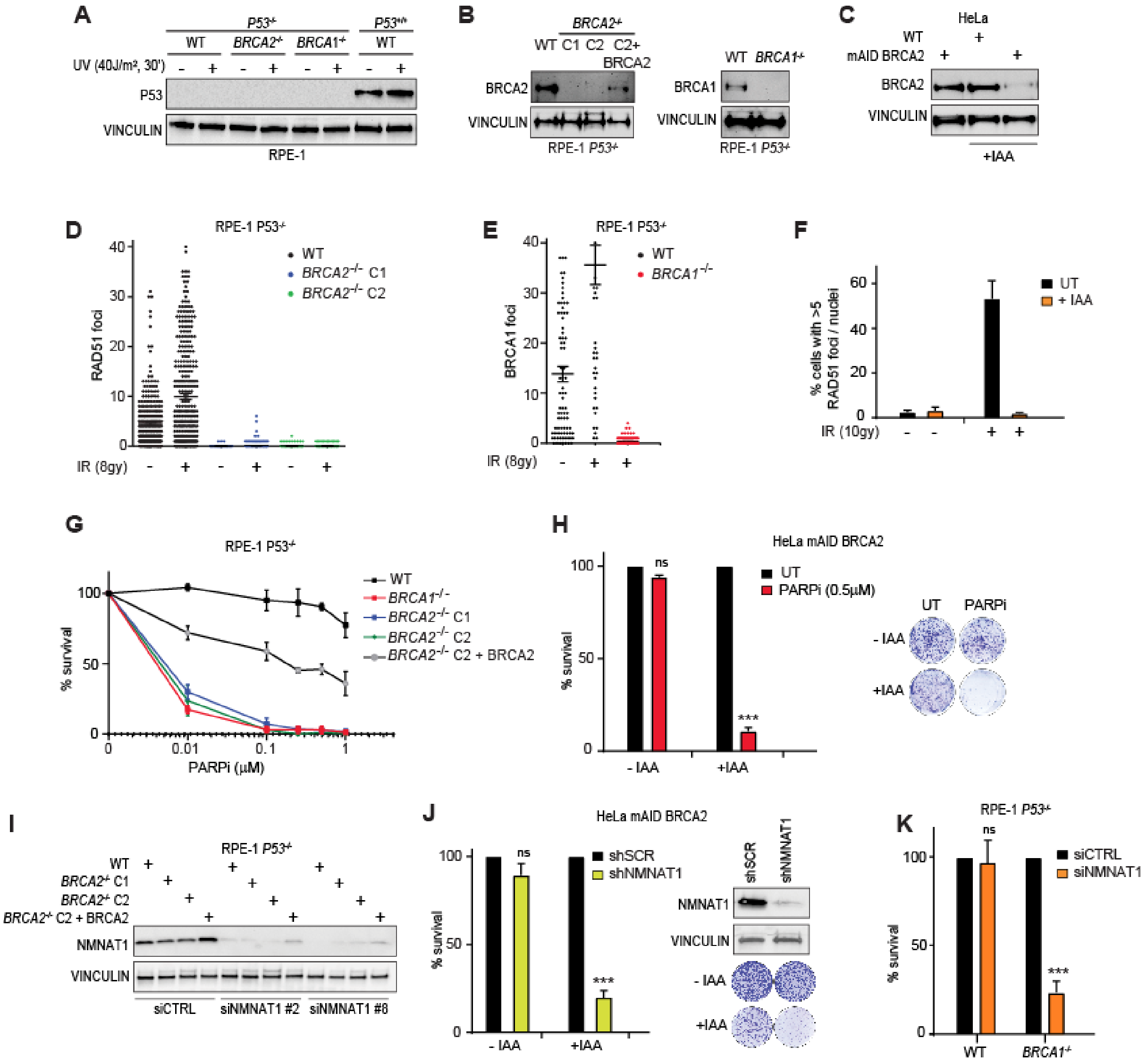

A
P53-/-
P53+/+
WT
BRCA2-/-
BRCA1-/-
UV (40J/m², 30')
P53
VINCULIN
RPE-1
B
BRCA2-/-
WT C1 C2 C2+ BRCA2
BRCA2
VINCULIN
RPE-1 P53-/-
WT BRCA1-/-
BRCA1
C
HeLa
WT
mAID BRCA2
+IAA
D
RPE-1 P53-/-
RAD51 foci
IR (8gy)
E
BRCA1 foci
F
% cells with >5 RAD51 foci / nuclei
UT
+ IAA
IR (10gy)
G
% survival
PARPi (μM)
H
HeLa mAID BRCA2
PARPi (0.5μM)
ns
- IAA
PARPi
I
BRCA2-/- C2 + BRCA2
NMNAT1
siCTRL
siNMNAT1 #2
siNMNAT1 #8
J
shSCR
shNMNAT1
K
siCTRL
siNMNAT1

**Fig. S1, related to Figure 1. *NMNAT1* is synthetic lethal with *BRCA1/2*. (A)**, Immunoblot analysis of p53 in both parental *P53*$^{+/+}$ RPE-1 cells and the indicated *P53*$^{-/-}$ derived clones (WT, *BRCA1*$^{-/-}$ and *BRCA2*$^{-/-}$) after exposure or not to UV light as indicated. **(B)**, Immunoblot analysis of BRCA2 (left *panel*) and BRCA1 (*right panel*) in both parental *P53*$^{-/-}$ RPE-1 cells and the *BRCA2*$^{-/-}$ or *BRCA1*$^{-/-}$ derived clones, respectively. C1 and C2 are two different *BRCA2*$^{-/-}$ clones (see methods section). C2+BRCA2 indicates clone C2 after complementation with BRCA2 full length cDNA. **(C)**, Immunoblot analysis of BRCA2 in both parental and mAID BRCA2 HeLa cells after treatment or not with auxin (IAA) at 500 nM for 16 hours. **(D)**, Quantification of RAD51 foci per nuclei in both parental RPE-1 cells and the *BRCA2*$^{-/-}$ derived clones C1 and C2 at 6 hours after treatment with ionizing radiation (IR, 8 Gy). **(E)**, Quantification of BRCA1 foci per nuclei in both parental *P53*$^{-/-}$ RPE-1 and *BRCA1*$^{-/-}$ cells at 6 hours after treatment with ionizing radiation (IR, 8 Gy). **(F)**, Quantification of RAD51 foci per nuclei in both parental and mAID BRCA2 HeLa cells after treatment or not with auxin (IAA) at 500 nM for 16 hours. Results are shown as percentage of cells with more than 5 RAD51 foci per nuclei after treatment or not with ionizing radiation (IR, 10 Gy). **(G)**, Clonogenic survival assay of RPE-1 cells as in **(B)** exposed to the indicated doses of the PARP inhibitor (PARPi) rucaparib. **(H)**, Quantification (*left panel*) and representative images (*right panel*) of clonogenic survival assay of mAID BRCA2 HeLa cells exposed to the indicated dose of the PARP inhibitor (PARPi) rucaparib in the presence or absence of 500 nM auxin (IAA). **(I),** Immunoblot analysis of NMNAT1 in RPE-1 cells as in (**B**, left *panel*) 72 hours after transfection with two different siNMNAT1 (siNMNAT1 #2 and #8) or siControl (siCTRL). **(J)**, Survival assay of mAID BRCA2 HeLa cells exposed to 500 nM auxin (IAA) 72 hours after transduction with lentiviral particles carrying either a shRNA targeting NMNAT1 (shNMNAT1) or a control shRNA (shSCR). The small inset contains the immunoblot analysis of

NMNAT1 in the same cells 72 hours after transduction. **(K)**, Survival assay of parental and *BRCA1*$^{-/-}$ RPE-1 cells as in (**B**, right panel) after transfection with either siNMNAT1 #2 or siControl (siCTRL). Vinculin is shown as a loading control in **(A-C )** and **(I)**. Data in **(H,J and K)** represent mean ± s.e.m. $n \geq 3$ (independent experiments); ns, non significant; ***, $p < 0.001$ (two-tailed *t* test).

**Fig. S2.**

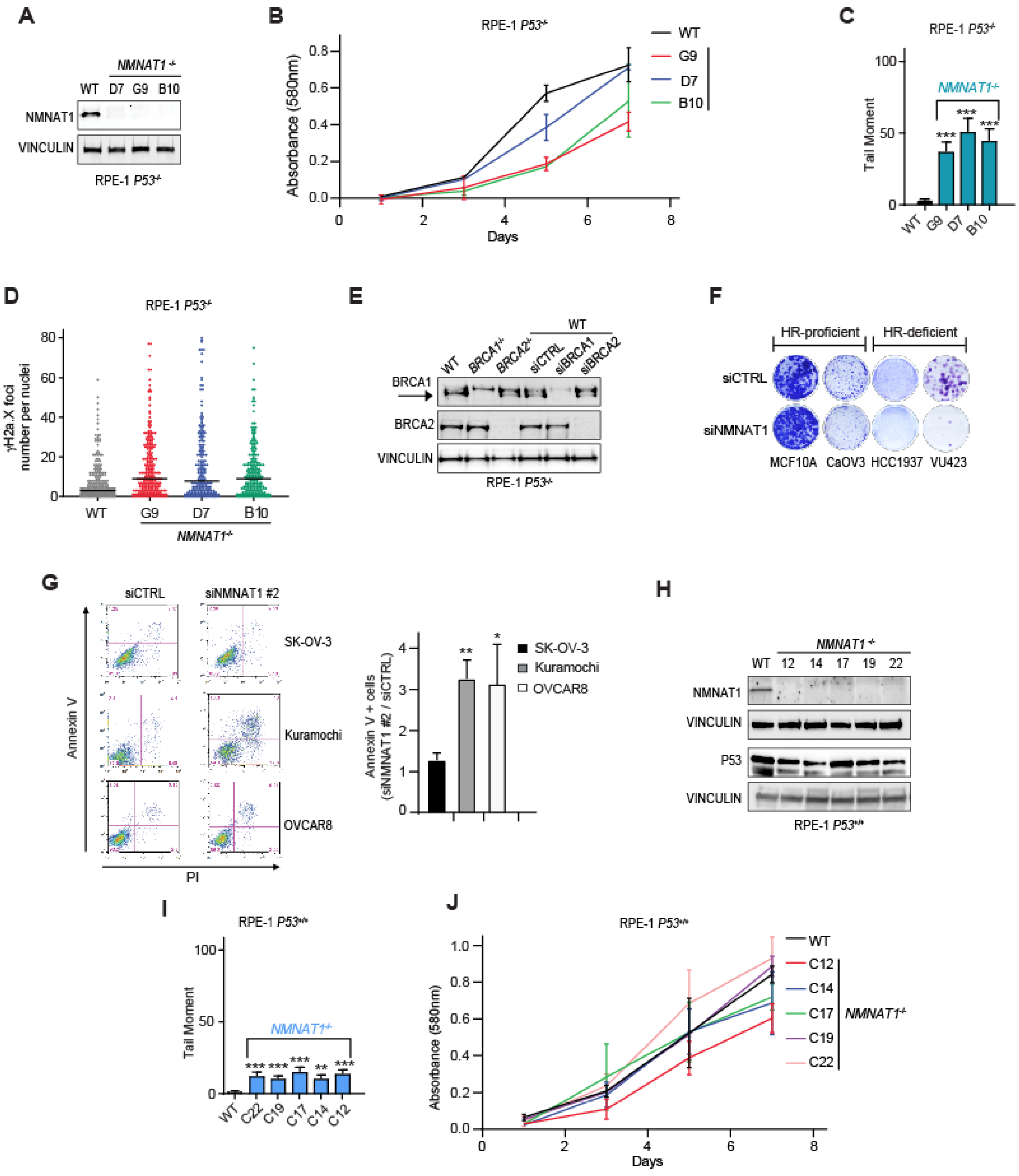
A
NMNAT1-/-
WT D7 G9 B10
NMNAT1
VINCULIN
RPE-1 P53-/-
B
RPE-1 P53-/-
Absorbance (580nm)
Days
WT
G9
D7
B10
C
RPE-1 P53-/-
Tail Moment
NMNAT1-/-
WT G9 D7 B10
D
RPE-1 P53-/-
γH2a.X foci number per nuclei
WT G9 D7 B10
NMNAT1-/-
E
WT BRCA1-/- BRCA2-/- siCTRL siBRCA1 siBRCA2
BRCA1
BRCA2
VINCULIN
RPE-1 P53-/-
F
HR-proficient
HR-deficient
siCTRL
siNMNAT1
MCF10A CaOV3 HCC1937 VU423
G
siCTRL
siNMNAT1 #2
SK-OV-3
Kuramochi
OVCAR8
Annexin V
PI
Annexin V + cells (siNMNAT1 #2 / siCTRL)
H
NMNAT1-/-
WT 12 14 17 19 22
NMNAT1
VINCULIN
P53
VINCULIN
RPE-1 P53+/+
I
RPE-1 P53+/+
Tail Moment
NMNAT1-/-
WT C22 C19 C17 C14 C12
J
RPE-1 P53+/+
Absorbance (580nm)
Days
WT
C12
C14
C17
C19
C22
NMNAT1-/-

**Fig. S2, related to Figure 1. *NMNAT1* is synthetic lethal with *BRCA1/2*. (A)**, Immunoblot analysis of NMNAT1 in both parental *P53*$^{-/-}$ RPE-1 cells and the indicated *NMNAT1*$^{-/-}$ derived clones (D7, G9 and B10). **(B)**, Proliferation assay of the cells as in **(A)** over one week. **(C and D)**, DNA damage quantification by alkaline COMET assay **(C)** or γH2a.x immunofluorescence **(D)** of the cells as in **(A)**, expressed as tail moment and γH2a.x foci number per nuclei respectively. **(E)**, Immunoblot analysis of BRCA1 and BRCA2 in *P53*$^{-/-}$ RPE-1 cells 72 hours following transfection with either *BRCA1* or *BRCA2* or control (siCTRL) siRNAs. *BRCA1*$^{-/-}$ and *BRCA2*$^{-/-}$ RPE-1 clones were used to confirm the specificity of the used antibodies. The arrow indicates the band corresponding to BRCA1 protein. **(F)**, Representative images of the survival assay of the indicated cell lines as quantified in (Fig. 1D) following transfection with siNMNAT1 #2 or siControl (siCTRL). **(G)**, Representative images and quantification of the apoptosis assay of the indicated cell lines 72 hours following transfection with siNMNAT1 #2 or siControl (siCTRL). Annexin-V positive cells are quantified as apoptotic cells and expressed as ratio between siNMNAT1 #2 to siCTRL in each cell line. **(H)**, Immunoblot analysis of NMNAT1 and p53 in both parental *P53*$^{+/+}$ RPE-1 cells and the indicated *NMNAT1*$^{-/-}$ derived clones (C12, C14, C17, C19 and C22). **(I and J)**, DNA damage quantification by alkaline COMET assay **(I)** and proliferation assay over one week **(J)** of the cells as in **(H)**. All data represent mean ± s.e.m. $n \geq 3$ (independent experiments); ns, non significant; *, $p < 0.05$; **, $p < 0.01$; ***, $p < 0.001$ (two-tailed *t* test).

**Fig. S3.**

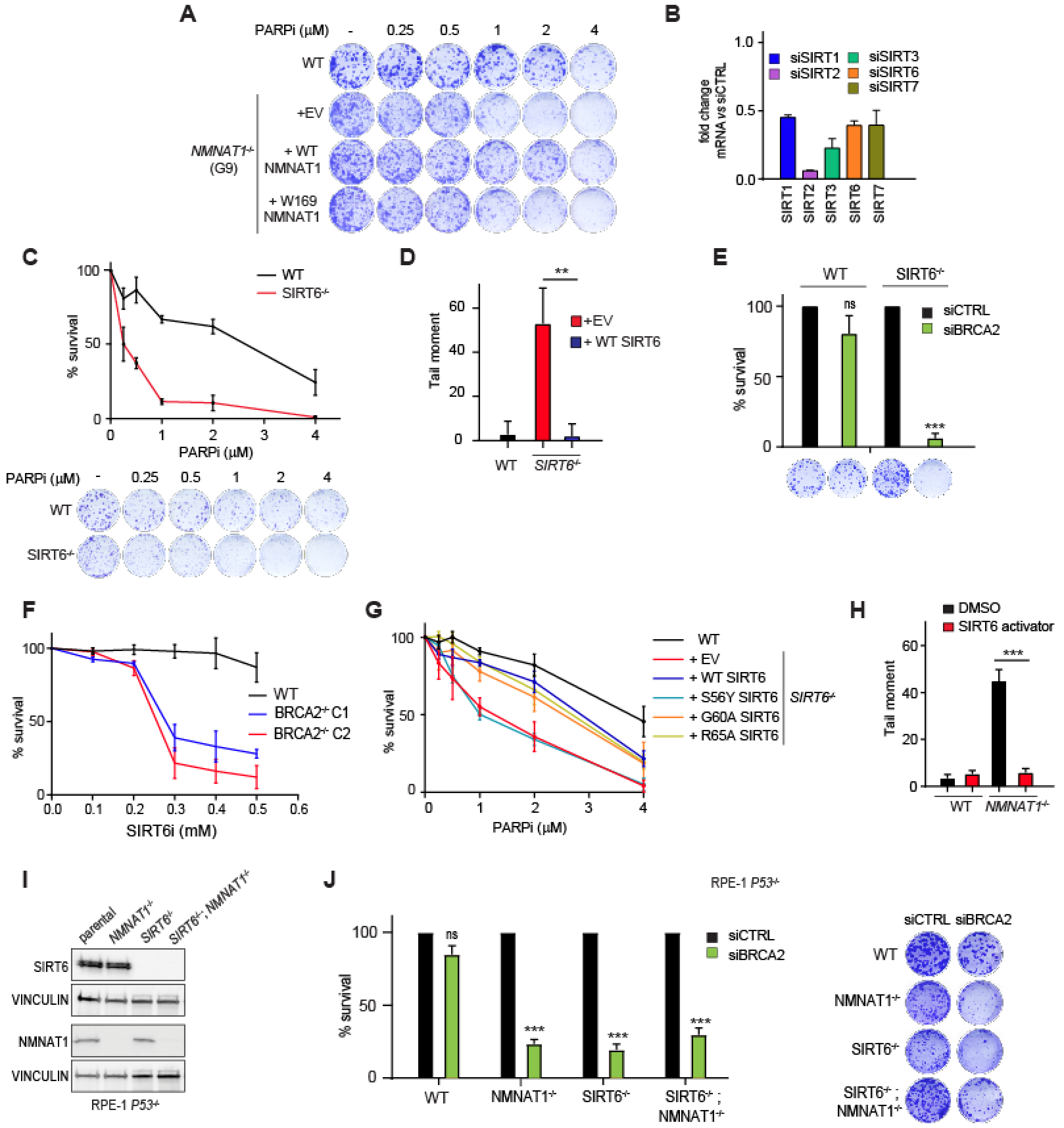
A
PARPi (µM)
-
0.25
0.5
1
2
4
WT
+EV
NMNAT1-/- (G9)
+ WT NMNAT1
+ W169 NMNAT1
B
siSIRT1
siSIRT2
siSIRT3
siSIRT6
siSIRT7
fold change mRNA vs siCTRL
SIRT1
SIRT2
SIRT3
SIRT6
SIRT7
C
WT
SIRT6-/-
% survival
PARPi (µM)
D
Tail moment
+EV
+ WT SIRT6
WT
SIRT6-/-
E
WT
SIRT6-/-
siCTRL
siBRCA2
% survival
ns
F
% survival
WT
BRCA2-/- C1
BRCA2-/- C2
SIRT6i (mM)
G
% survival
WT
+ EV
+ WT SIRT6
+ S56Y SIRT6
+ G60A SIRT6
+ R65A SIRT6
SIRT6-/-
PARPi (µM)
H
DMSO
SIRT6 activator
Tail moment
WT
NMNAT1-/-
I
parental
NMNAT1-/-
SIRT6-/-
SIRT6-/-, NMNAT1-/-
SIRT6
VINCULIN
NMNAT1
VINCULIN
RPE-1 P53-/-
J
RPE-1 P53-/-
% survival
siCTRL
siBRCA2
ns
WT
NMNAT1-/-
SIRT6-/-
SIRT6-/- ; NMNAT1-/-

**Fig. S3, related to Figure 2. SIRT6 functions downstream of NMNAT1 in the survival of HRD cells. (A)**, Representative images of the clonogenic survival assay shown in Fig. 2C. **(B)**, Quantification by qPCR of the mRNA levels of the indicated nuclear sirtuins in *P53*$^{-/-}$ RPE-1 cells 72 hours after transfection with either sirtuin (siSIRT) or control (siCTRL) siRNAs. Data are normalized to housekeeping gene (RPLP01) expression and mRNA level for each analyzed sirtuin is expressed as fold change between the indicated siSIRT sample over the control (siSIRT/siCTRL). **(C)**, Survival assay of *SIRT6*$^{-/-}$ and parental *P53*$^{-/-}$ RPE-1 cells exposed to the indicated doses of the PARP inhibitor (PARPi) rucaparib. **(D)**, DNA damage quantification by alkaline COMET assay of parental and *SIRT6*$^{-/-}$ *P53*$^{-/-}$ RPE-1 cells complemented (+ WT SIRT6) or not (empty vector, EV) with SIRT6 cDNA. **(E)**, Quantification (*upper panel*) and representative images (*lower panel*) of the clonogenic formation of *SIRT6*$^{-/-}$ and parental *P53*$^{-/-}$ RPE-1 cells following transfection with either *BRCA2* or control (siCTRL) siRNAs. **(F),** Quantification of clonogenic survival assay of wild type (WT) and *BRCA2*$^{-/-}$ (clones C1 and C2) RPE-1 cells in response to the indicated doses of the SIRT6 inhibitor OSS128167 (SIRT6i). **(G)**, Quantification of clonogenic survival assay of parental (WT) and *SIRT6*$^{-/-}$ *P53*$^{-/-}$ RPE-1 cells complemented or not (empty vector, EV) with the indicated SIRT6 cDNAs in response to the indicated doses of the PARP1 inhibitor rucaparib (PARPi). **(H)**, DNA breaks quantification by alkaline COMET assay of parental (WT) and *NMNAT1*$^{-/-}$ *P53*$^{-/-}$ RPE-1 cells treated or not (DMSO) for 3 days with 10 μM of the SIRT6 activator MDL800. **(I)**, Immunoblot analysis of NMNAT1 and SIRT6 in parental *P53*$^{-/-}$ RPE-1 cells and the *NMNAT1*$^{-/-}$ (G9), *SIRT6*$^{-/-}$ and *SIRT6*$^{-/-}$ *NMNAT1*$^{-/-}$ derived clones. **(J)**, Quantification of clonogenic survival assay (*left panel*) and representative images (*right panel*) of RPE-1 cells as in **(I)** after transfection with si*BRCA2* or control (siCTRL). All data represent mean

± s.e.m. $n \geq 3$ (independent experiments); ns, non significant; **, $p < 0.01$; ***, $p < 0.001$ (two-tailed $t$ test).

### Fig. S4

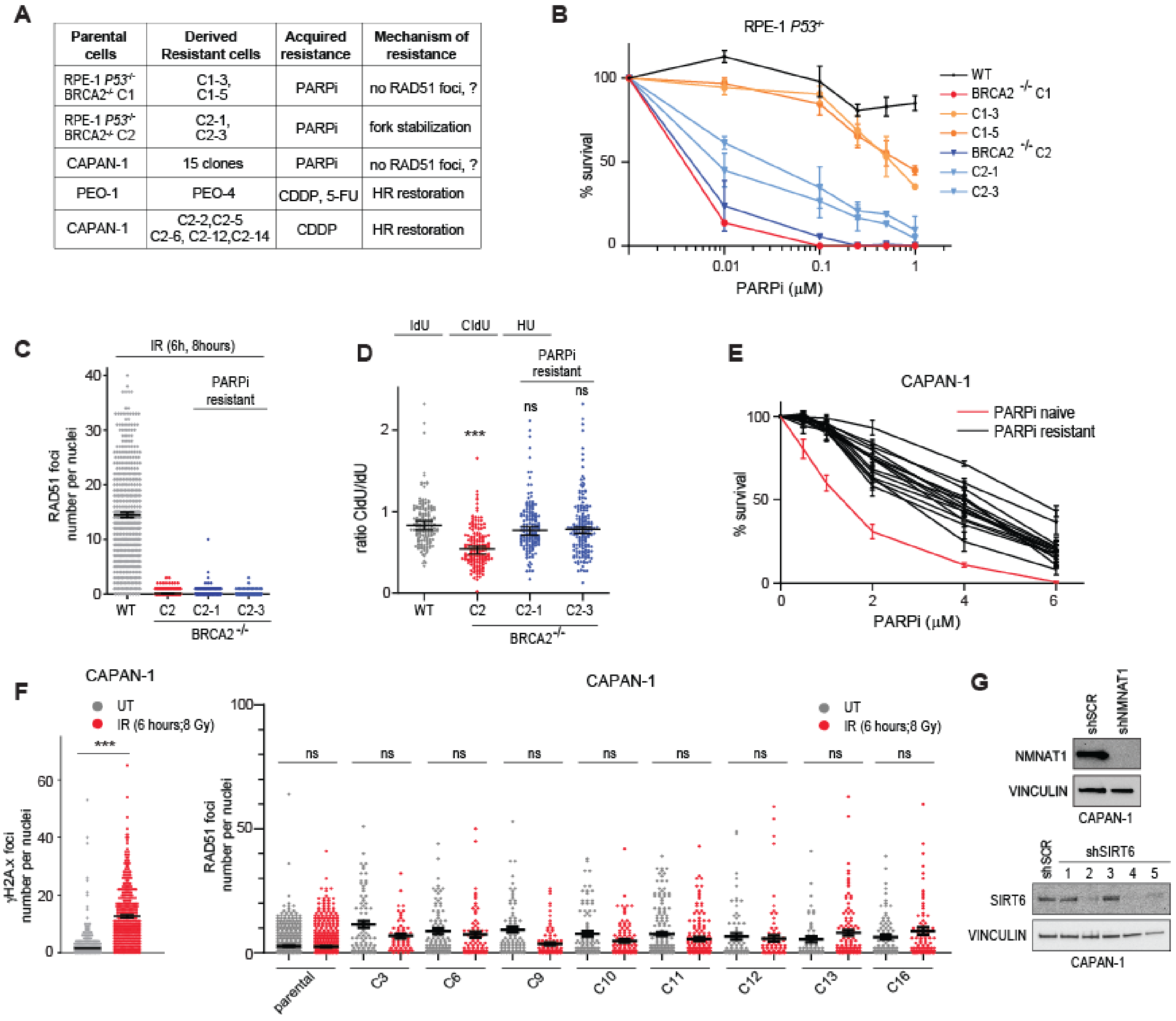
A
| Parental cells | Derived Resistant cells | Acquired resistance | Mechanism of resistance |
| --- | --- | --- | --- |
| RPE-1 P53-/- BRCA2-/- C1 | C1-3, C1-5 | PARPi | no RAD51 foci, ? |
| RPE-1 P53-/- BRCA2-/- C2 | C2-1, C2-3 | PARPi | fork stabilization |
| CAPAN-1 | 15 clones | PARPi | no RAD51 foci, ? |
| PEO-1 | PEO-4 | CDDP, 5-FU | HR restoration |
| CAPAN-1 | C2-2,C2-5 C2-6, C2-12,C2-14 | CDDP | HR restoration |

B
RPE-1 P53-/-
% survival
PARPi (µM)
WT
BRCA2 -/- C1
C1-3
C1-5
BRCA2 -/- C2
C2-1
C2-3
C
IR (6h, 8hours)
PARPi resistant
RAD51 foci number per nuclei
WT
C2
C2-1
C2-3
BRCA2 -/-
D
IdU
CldU
HU
PARPi resistant
ratio CldU/IdU
***
ns
E
CAPAN-1
PARPi naive
PARPi resistant
% survival
PARPi (µM)
F
CAPAN-1
UT
IR (6 hours;8 Gy)
γH2A.x foci number per nuclei
RAD51 foci number per nuclei
parental
C3
C6
C9
C10
C11
C12
C13
C16
G
shSCR
shNMNAT1
NMNAT1
VINCULIN
CAPAN-1
shSIRT6
SIRT6

**Fig. S4, related to Figure 3. Inhibition of NMNAT1 or SIRT6 kills PARPi-resistant HRD cells, regardless the mechanism of resistance. (A)**, Schematic of the drug-resistant cell lines used in this work. The mechanism of acquired drug resistance is also indicated. **(B)**, Clonogenic formation of wild type (WT), *BRCA2*$^{-/-}$ (clones C1 and C2) and the indicated *BRCA2*$^{-/-}$-derived PARPi-resistant RPE-1 cells (C1-3, C1-5, C2-1 and C2-3) following exposure to the indicated doses of the PARP1 inhibitor (PARPi) rucaparib. **(C)**, Quantification of RAD51 foci per nuclei in wild type (WT), *BRCA2*$^{-/-}$ (clone C2) and the indicated clone C2-derived PARPi-resistant (C2-1 and C2-3) RPE-1 cells 8 hours after exposure to ionizing radiation (6 Gy). **(D)**, Quantification of replication fork degradation as measured by combing assay of the cells as in **(C)**. Rate of fork degradation is expressed as ratio in terms of tract length between CIdU and IdU incorporated fibers after a 3 hours treatment with hydroxyurea (HU, 4 mM). **(E)**, Clonogenic formation of parental (naive) and derived PARPi-resistant CAPAN-1 cells following exposure to the indicated doses of the PARP inhibitor (PARPi) rucaparib. **(F)**, Quantification of the RAD51 foci per nuclei (*right panel*) in cells as in **(E)** 6 hours after exposure (IR) or not (UT) to 8 Gy of ionizing radiation. *Left panel* shows the quantification of γH2a.x foci formation after treatment of parental CAPAN-1 cells as in *right panel*. **(G)**, Immunoblot analysis of NMNAT1 (*upper panel*) and SIRT6 (*lower panel*) in CAPAN-1 cells after transduction with shNMNAT1 (*upper panel*) or shSIRT6 (*lower panel*) lentiviral particles. Lentiviral particles carrying control shRNA (shSCR) were used as control. Five different shRNA have been tested for SIRT6 knockdown efficiency and SIRT6 #4 has been used in the other experiments of this manuscript. All data represent mean ± s.e.m. $n \geq 3$ (independent experiments); ns, non significant; ***, $p < 0.001$ (two-tailed *t* test).

**Fig. S5**

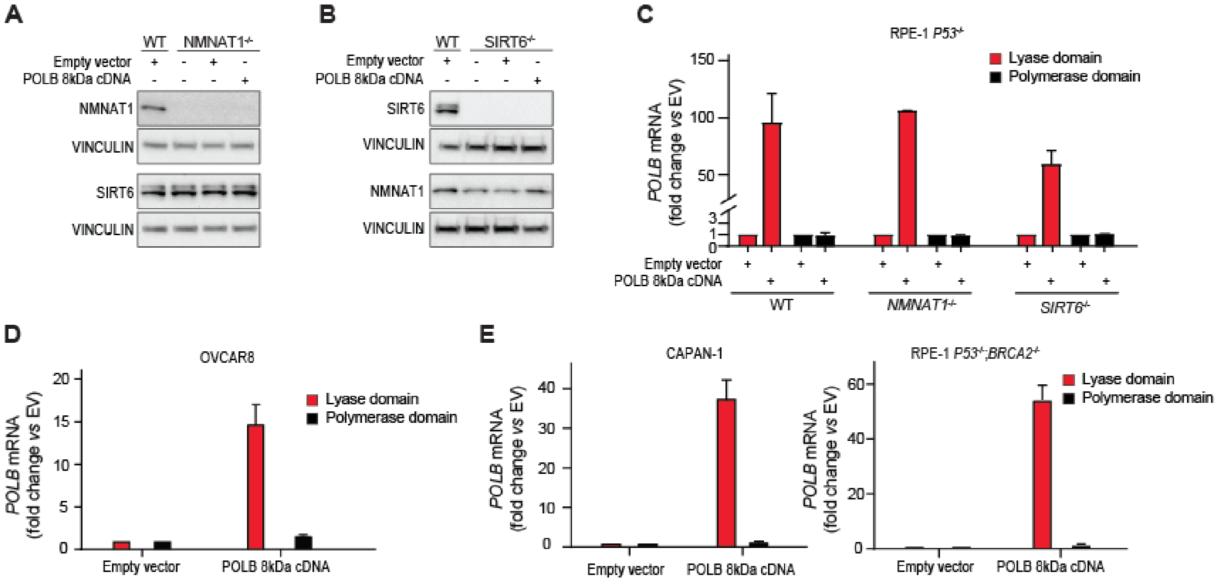


**Fig. S5, related to Figure 4. Over-expression of POLB lyase domain overcomes cytotoxicity of NMNAT1 or SIRT6 inhibition in HRD cells. (A and B),** Immunoblot analysis of NMNAT1 and SIRT6 following over-expression of POLB 8kDa lyase domain in *NMNAT1*$^{-/-}$ (G9) **(A)** and *SIRT6*$^{-/-}$ **(B)** *P53*$^{-/-}$ RPE-1 clones. **(C-F)**, Quantification by qPCR of the mRNA levels of both polymerase and lyase POLB domains following over-expression or not (EV) of the POLB 8kDa lyase domain (POLB 8kDa cDNA) in *NMNAT1*$^{-/-}$ *P53*$^{-/-}$ (G9) and *SIRT6*$^{-/-}$ *P53*$^{-/-}$ RPE-1 **(C)**, OVCAR8 **(D)**, CAPAN-1 **(E)** and *BRCA2*$^{-/-}$ *P53*$^{-/-}$ RPE-1 clone 2 (C2) **(F)**. Data are normalized to housekeeping gene (RPLP01) expression and mRNA level for each POLB domain is expressed as fold change over the control (EV). All data represent mean ± s.e.m. $n \geq 3$ (independent experiments).
